# Causal coupling between neural activity, metabolism, and behavior across the *Drosophila* brain

**DOI:** 10.1101/2020.03.18.997742

**Authors:** Kevin Mann, Stephane Deny, Surya Ganguli, Thomas R. Clandinin

## Abstract

Coordinated activity across networks of neurons is a hallmark of both resting and active behavioral states in many species, including worms, flies, fish, mice and humans^1–5^. These global patterns alter energy metabolism in the brain over seconds to hours, making oxygen consumption and glucose uptake widely used proxies of neural activity^6,7^. However, whether changes in neural activity are causally related to changes in metabolic flux in intact circuits on the sub-second timescales associated with behavior, is unknown. Moreover, it is unclear whether transitions between rest and action are associated with spatiotemporally structured changes in neuronal energy metabolism. Here, we combine two-photon microscopy of the entire fruit fly brain with sensors that allow simultaneous measurements of neural activity and metabolic flux, across both resting and active behavioral states. We demonstrate that neural activity drives changes in metabolic flux, creating a tight coupling between these signals that can be measured across large-scale brain networks. Further, these studies reveal that the initiation of even minimal behavioral movements causes large-scale changes in the pattern of neural activity and energy metabolism, revealing unexpected structure in the functional architecture of the brain. The relationship between neural activity and energy metabolism is likely evolutionarily ancient. Thus, these studies provide a critical foundation for using metabolic proxies to capture changes in neural activity and reveal that even minimal behavioral movements are associated with changes in large-scale brain network activity.

---

Three technologies have been widely used to measure changes in neural activity across whole brain volumes. In humans, non-human primates, and rodents, functional magnetic resonance imaging (fMRI) uses blood oxygen level dependent (BOLD) signals to capture changes in oxygenated blood flow as a proxy for neural activity, with a temporal resolution of seconds and a spatial resolution of millimeters^6^. Fluorodeoxyglucose positron emission tomography (FDG-PET) captures changes in glucose uptake, with a temporal resolution of tens of seconds and a typical spatial resolution of centimeters^7,8^. Simultaneous imaging methods in humans have demonstrated that FDG-PET intrinsic (i.e., task-free) brain networks spatially overlap with BOLD networks, suggesting a relationship between glucose uptake and blood oxygenation^9^. In parallel, in genetic model organisms, imaging approaches that use fluorescent sensors to measure changes in intracellular calcium concentrations have been widely applied to capture neural activity with single cell resolution across large brain areas^2–4,10–13^. However, none of these approaches have allowed direct, simultaneous, brain-wide intracellular measurements of changes in both neural activity and metabolic flux.

Understanding brain function on mesoscale often relies on measuring functional connectivity. Functional connectivity networks are defined by correlated changes in neural activity between regions over time, where the strength of each connection in the network is defined by the magnitude of the correlation between the activity patterns in pairs of regions^5^. These correlations capture large scale interactions across the brain that reflect brain regions coordinating their activity to shape behavior. Such network structures have been derived using BOLD, FDG-PET, and fluorescent calcium sensors in a variety of species^2,3,5,9^. However, no temporally coherent, direct measure of both metabolic flux and neural activity has been achieved at the mesoscale to measure network activity either at rest or during behavior.

In *Drosophila*, neural activity has been measured across the whole brain using fluorescent calcium and voltage sensors, revealing correlated patterns of activity between specific brain regions (Fig. 1a;)^3,13^. In this approach, registering neural activity signals to a common anatomical atlas allows signals from anatomically defined regions to be compared across animals^4^. These tools, combined with the recent development of optical sensors for metabolic flux and red-shifted calcium indicators^14–17^, allow for the simultaneous imaging of both metabolism and neural activity across the entire brain of the fly. Here, we utilize these sensors to determine the spatiotemporal relationship between metabolic flux and neural activity.

**Figure 1.**
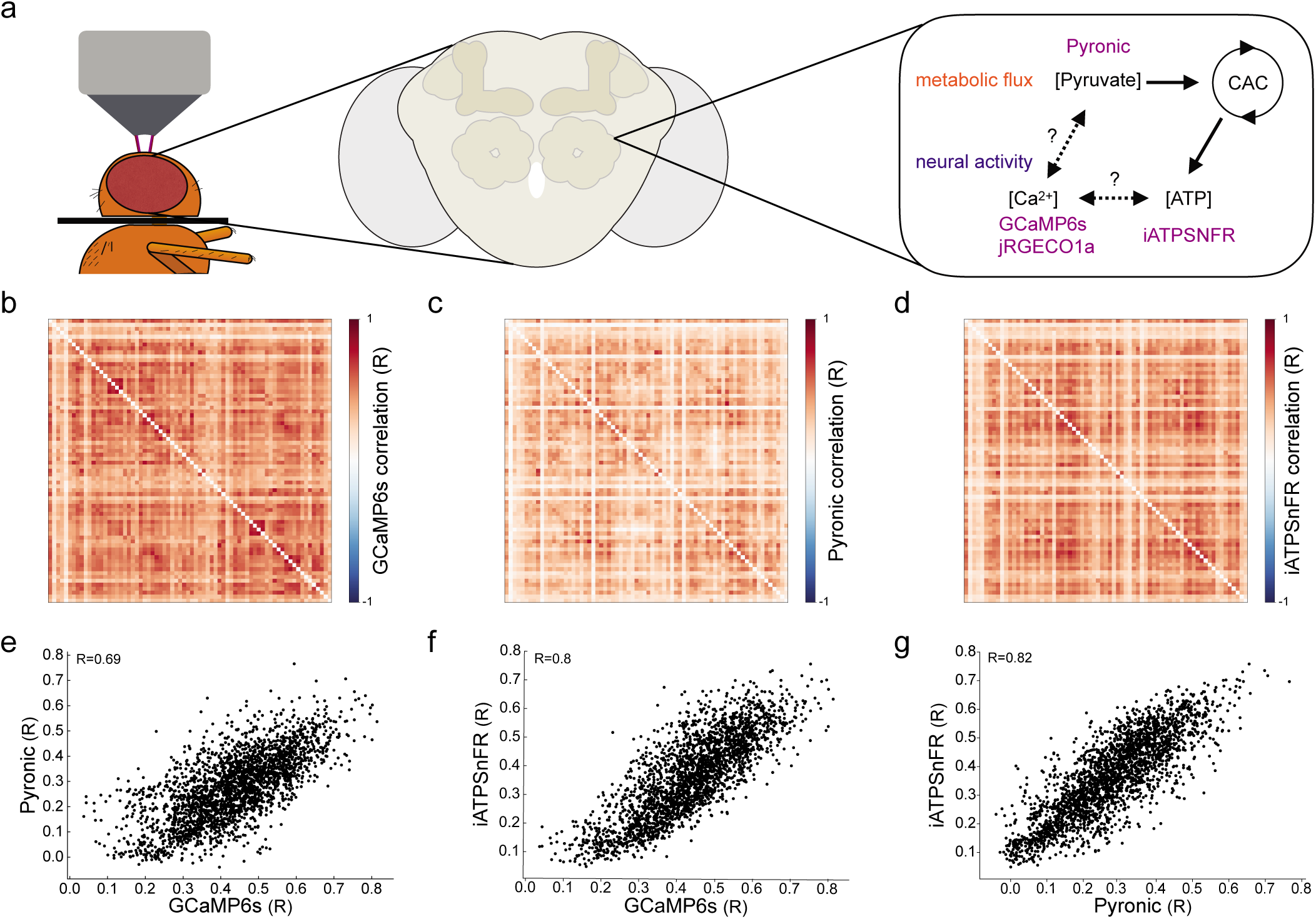
Metabolic and neural networks are highly correlated across the brain. (a). (left) Schematic illustration of the preparation that allows 2-photon imaging across the fly brain. (middle) Cartoon of the imaged region of the fly brain. (right) Schematic illustration of a neuronal process, denoting the metabolic pathways that lead to ATP production, and the sensors that were used to measure changes in intracellular calcium concentration (GCaMP6s, jrGECO1a), pyruvate concentration (Pyronic) and ATP concentration (iATPSnFR). (b-d). Matrices of pairwise correlations between brain regions. (b). GCaMP6s. (c). Pyronic. (d). iATPSnFR. (e-g) Scatter plots of the pairwise correlations between matrices. (e). Pyronic vs. GCaMP6s. (f). iATPSnFR vs GCaMP6s. (g). iATPSnFR vs Pyronic. n = 12 flies for GCaMP6s, n = 10 for Pyronic, n = 10 for iATPSNFr.

## Neural activity and metabolic flux are correlated across the brain

Correspondence between neural activity and metabolism can be measured using genetically encoded sensors, combined with brain-wide imaging, in immobilized animals (Fig. 1a)^3^. We hypothesized that if normal variations in neural activity in the intact brain were closely coupled to variations in intracellular energy production, a functional connectivity network could be detected using a sensor that measures changes in energy metabolism. To do this, we took advantage of a FRET-based sensor, Pyronic, whose output responds rapidly to changes in intracellular pyruvate concentration (Fig. 1a)^14,15^, as well as iATPSnFR, a circularly permuted GFP sensor of changes in ATP concentration^16^, and compared these signals to intracellular calcium levels measured using GCaMP6s^18^. As changes in both the citric acid cycle (CAC) and glycolysis (as well as other metabolic pathways) alter pyruvate flux and ATP levels, we reasoned that changes in Pyronic and iATPSnFR signals should probe whether changes in metabolic activity are correlated within and between brain regions.

We adapted methods previously developed for whole-brain calcium imaging, expressing both of these metabolic sensors pan-neuronally, along with a structural marker tdTomato, and imaged the entire central brains of immobilized animals^3^, see Methods. Signals were continuously measured for approximately 20 minutes at 1.2Hz (at an isotropic spatial resolution of 3um), before acquisition of a high-resolution structural scan using tdTomato (at an isotropic spatial resolution of 1um). Next, each brain was aligned with a template brain^19^, and a standard atlas was used to extract each of these signals from 60 anatomically defined regions^20^. In each brain region, all of these signals varied continuously, with no obvious periodicity. Next, we correlated the time series of each signal from each pair of regions, and observed strong correlations between some pairs, but not others (Extended Data Fig. 1). To assess whether these correlations were stereotyped between animals, we then compared the average correlations between all pairs of regions, across all flies, providing us with a functional connectivity map of metabolic flux (Fig. 1b-d, Extended Data Fig. 2). Remarkably, all three connectivity maps were highly structured, demonstrating that metabolic flux at the level of pyruvate and ATP levels, as well as neural activity, are coordinated between specific regions of the brain in a manner that is highly stereotyped from animal to animal.

We next tested whether there were similarities between each of the metabolic flux networks and the calcium activity network. To do this, we computed the pairwise correlations between each correlation matrix (Fig. 1e-g). These comparisons revealed that these correlation matrices were all very similar (R=0.69 for Pyruvate versus Calcium, R=0.80 for ATP versus Calcium, and R=0.82 for ATP versus Pyruvate), even though the metabolic flux and neural activity measures were made in different animals, using different sensors, targeting different aspects of energy metabolism. As the connectivity patterns were remarkably similar in different animals, we developed a procedure to investigate the link between metabolism and neural activity in the same animal. Specifically, we co-expressed both Pyronic and the red-shifted calcium indicator jRGECO1a pan-neuronally, and simultaneously measured both metabolic flux and neural activity (n=24 animals). Under these conditions, Pyronic and jRGECO1a connectivity matrices were also highly similar (Extended Data Fig. 3). Taken together, these data demonstrate that the functional connectivity structure of neural signals throughout the nervous system is mirrored in the corresponding structure of metabolic flux, suggesting an intimate link between neural activity and metabolism.

## Changes in neural activity drive changes in metabolic flux

We next sought to determine the relationship between metabolic flux and neural activity at the level of individual brain regions and voxels (Fig. 2). Qualitative and quantitative comparisons of the simultaneously recorded measurements of these signals revealed a significant correlation between the two signals, pooled across all frequency bands (Fig. 2a-c). Strikingly, these correlations were much stronger when the flux and neural activity signals were filtered to select for low frequencies, rather than high frequencies (Fig. 2d-g). Importantly, these correlations were eliminated by shuffle controls that swapped either temporal relationships or regional identities (Fig. 2h). Thus, low frequency changes in intracellular calcium levels, corresponding to the timescales of tens of seconds, are highly correlated with changes in metabolic flux.

**Figure 2.**
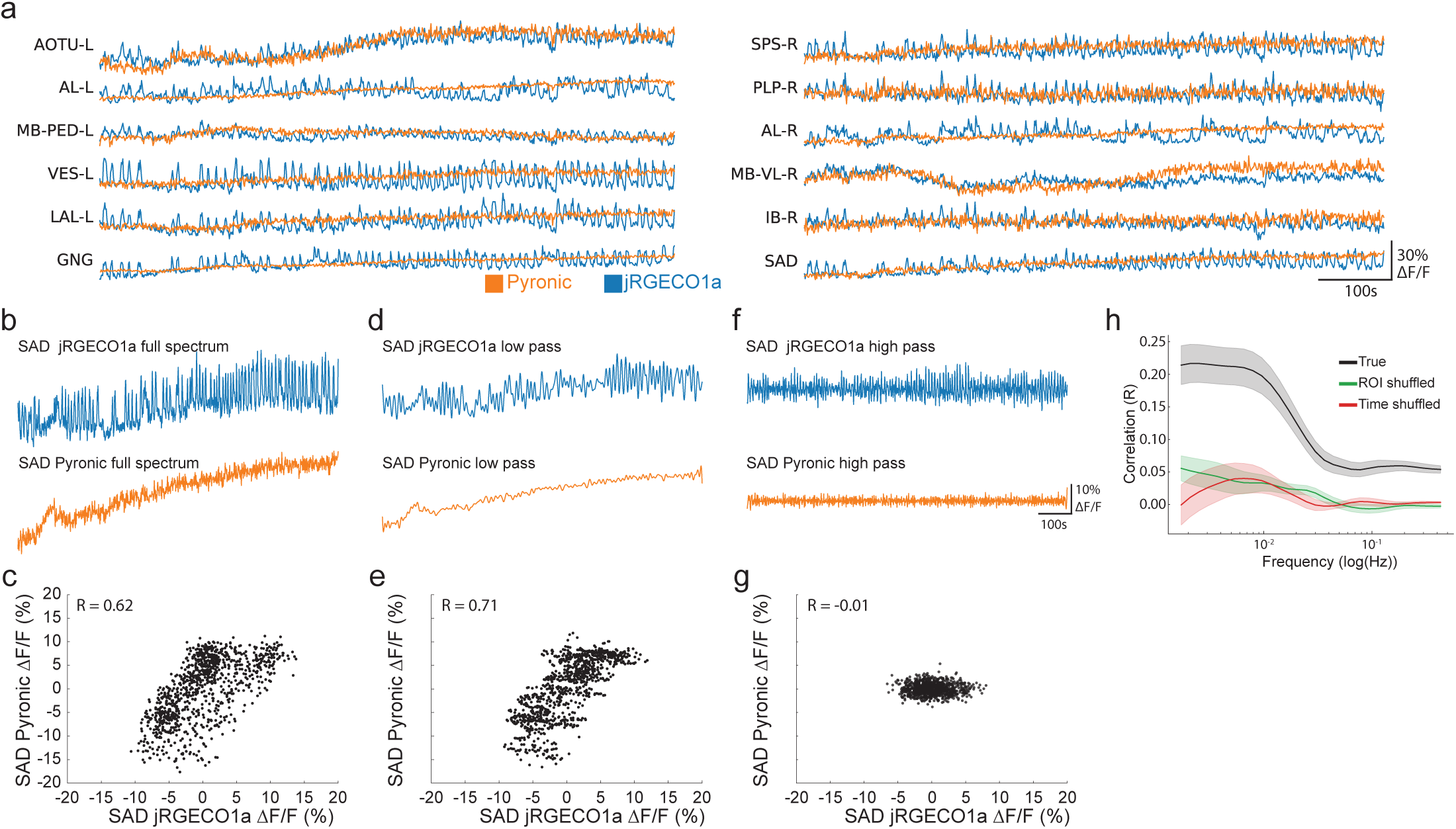
Simultaneous measurements of neural activity and metabolic flux reveal correlations dominated by low frequencies. (a). Traces displaying Pyronic signals (orange) and jRGECO1a signals (blue), across 12 different brain regions. (b-g) Comparison of jRGECO1a and Pyronic signals within a single brain region, the Saddle (SAD). (b). Traces of Pyronic and jRGECO1a signals including all frequency components. (c). Pairwise comparison of Pyronic and jRGECO1a signals including all frequency components and the correlation between these signals. (d, e). As in (b,c), but filtered to include only low frequency (< 0.1 Hz) components. (f,g). As in (b,c), but filtered to include only high frequency (> 0.1Hz) components. (h). Pairwise correlations between Pyronic and jRGECO1a signals measured in each brain region, as a function of frequency (black trace). Shuffle control in which pairwise comparisons were done between brain regions whose identities have been shuffled (green trace), or in which the signals have been shuffled in time (red trace) n = 24 animals, mean ± SEM (shading).

We next sought to examine the causal relationship underpinning the correlations between metabolic flux and neural activity. To do this, we imaged flies simultaneously expressing Pyronic and jRGECO1a before and after bath-application of tetrodotoxin (TTX). TTX blocks voltage-gated sodium channels, preventing action potential generation^21^, thereby strongly inhibiting neural activity (though activity in non-spiking neurons can persist). If changes in neural activity drive changes in metabolic flux, then blocking neural activity should disrupt both the neural and metabolic functional connectivity maps by eliminating correlations between these signals. Bath application of TTX dramatically reduced the fluctuations in both the jRGECO1a and Pyronic signals (Fig. 3a,b), suppressing signals across a wide range of frequencies (Fig. 3c). Moreover, blocking neural activity also largely eliminated the stereotyped correlations between these signals, across the entire brain (Fig. 3d, e). Thus, the observed metabolic network is substantially the product of neural activity.

**Figure 3.**
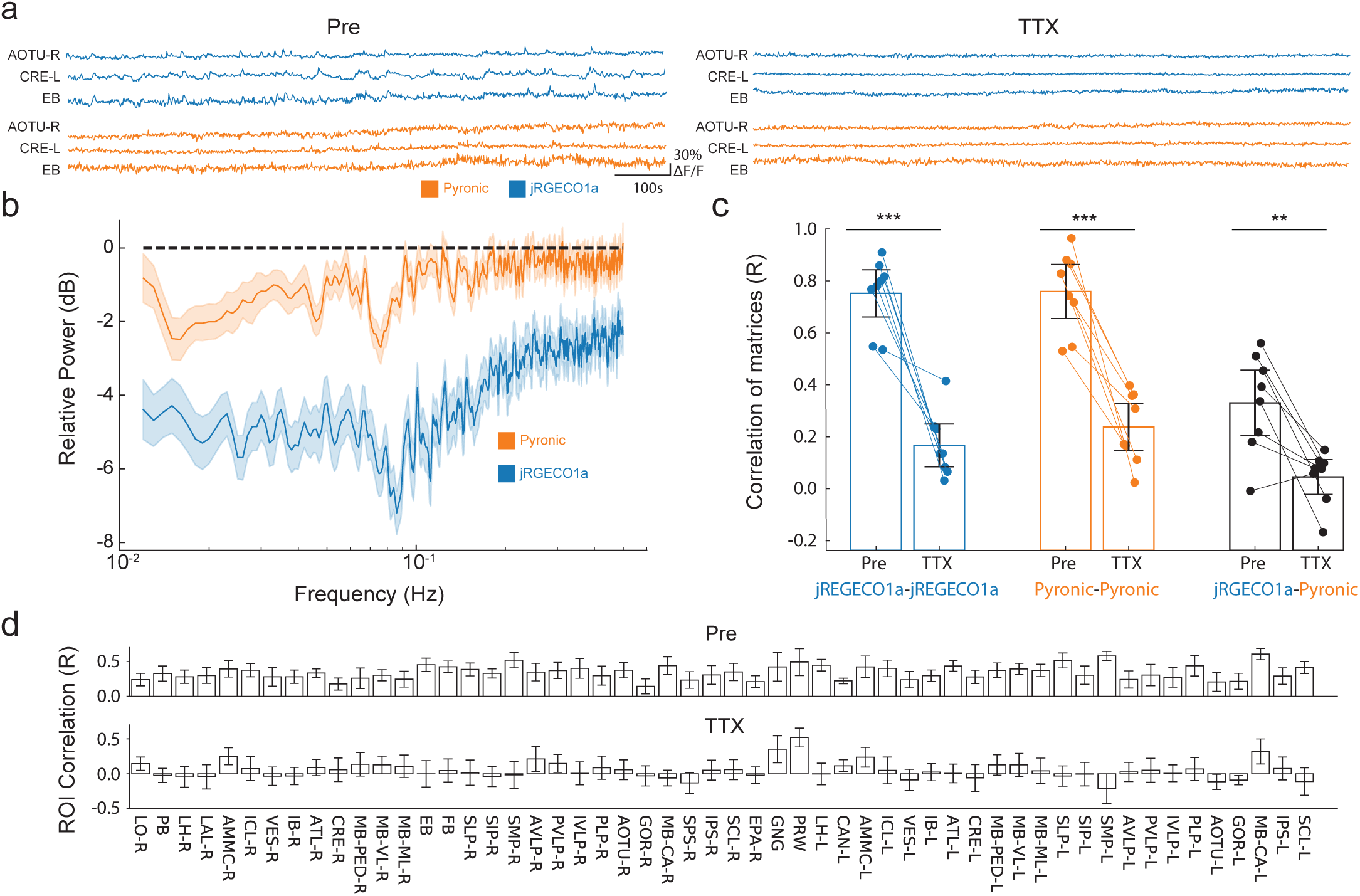
Neural activity drives metabolic flux in the brain. (a). jRGECO1a (blue) and Pyronic (orange) traces in three different brain regions before (left) and after (right) application of tetrodotoxin (TTX). (b). Comparison of the variance of the signals of each ROI before and after TTX application, n=8 flies, mean ± SEM for each region. (c). The relative reduction in signal power caused by TTX application, as a function of frequency across all brain regions and flies (n= 54 regions and 8 flies, mean ± 95% confidence interval (shading)). (d). Correlation of the correlation maps between flies, before and after TTX application, across all brain regions, for jRGECO1a (calcium, blue dots), Pyronic (pyruvate, orange dots), and for the correlations between jRGECO1a and Pyronic (black dots, n=8 flies, mean ± 95% confidence interval) *** P < P < 0.0004, ** P < 0.005. (e). Region by region correlations between calcium and pyronic signals, across all flies, before TTX application (upper row) and after TTX application (lower row), n=8 flies, mean ± SEM.

## Initiation of movement restructures both neural activity and metabolic flux networks

These studies demonstrate that changes in neural activity strongly shape metabolic flux. However, how these signals might be coordinately altered by the engagement of behavior remains unknown. To address this, we simultaneously imaged Pyronic and jRGECO1a, while recording leg movements to determine if the fly was active or quiescent (see Methods). We then trained a generalized linear model (GLM) using either neural activity or metabolic flux to predict bouts of movement (Fig. 4a-c and Extended Data Fig 4). Strikingly, changes in neural activity in specific, stereotyped regions of the brain were able to predict the timing of movement bouts, even when these bouts were very brief (∼1 second) (Fig. 4b,c and Extended Data Fig 4; R = 0.49-0.76). The accuracy of behavioral predictions spanned all but the lowest frequencies observed, closely tracking the power spectrum of behavior itself (Extended Data Fig. 5). Conversely, models that attempt to predict bouts of activity from metabolic flux performed relatively poorly, but still performed above chance (Fig. 4b,c and Extended Data Fig. 4; R = 0.11-0.25 across all flies). These correlations were highest at intermediate frequencies, consistent with both the power spectrum of behavior, and the SNR of Pyronic (Fig. 4c; Extended Data Fig. S5).

**Figure 4.**
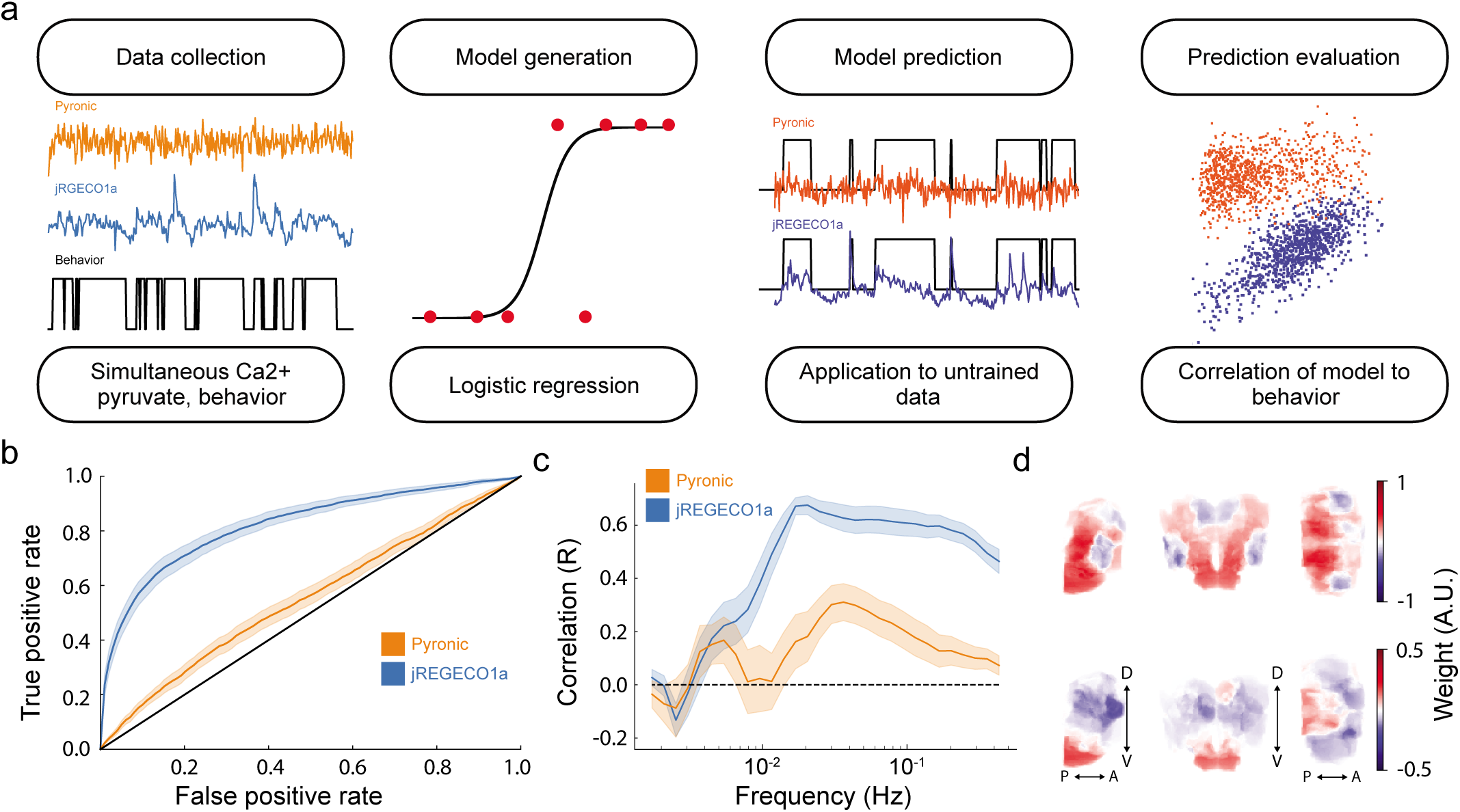
Neural activity and metabolic flux are correlated with behavior in specific regions. (a). Schematic of the data processing and analysis pipeline utilized. (*i*). Traces of Pyronic, jRGECO1a, and behavior (movement of the legs). (ii). Half of the data set was used to train a logistic regression model relating neural activity and metabolic flux to behavior. (iii). Predicted behavioral outputs were generated using the withheld data and were compared to the actual behavior during those time periods. (iv) Model prediction was evaluated by correlating predicted behavior to observed behavior. (b) Receiver operator curve (ROC) showing a good prediction of behavior across all flies, using models based on jRGECO1a (blue line, area under curve (AUC) = 0.82, P < 0.0001, one-tailed t-test against 0.5), and poorer but significant prediction of behavior using Pyronic (orange line, AUC = 0.54, P<0.05, on-tailed t-test against 0.5). (c). Comparisons of correlations between predictions of behavior based on jRGECO1a (blue line) and Pyronic (orange line) across a range of frequencies. Pyronic is a better predictor of behavior in the middle of the frequency range (0.04Hz), while jRGECO1a predicts behavior at those and higher frequencies (mean ± 95% confidence interval). (d). Average weights of each ROI generated from the logistic regression model when computed at the peak frequency of correlation for metabolic flux and behavior (0.04Hz). Images are sagittal, coronal, and axial views, respectively, of the central brain, and colored by weight (scale bar). n = 12 jRGECO1a, n = 8 Pyronic.

To probe the generality of the spatial structure of these GLMs across flies, we computed the average weights used in each model, for each brain region, at the optimal predictive frequency (Fig. 4c). Weights generated from calcium signals revealed a structured map of regions predictive of behavior (Fig. 4d). Weights generated from metabolic flux signals were also structured, capturing a subset of the most strongly weighted regions in the neural activity maps, and correlating with the overall structure observed (R = 0.36)(Fig. 4d). Thus, there is a region-specific pattern of common neural and metabolic load associated with behavior.

To better define the regions correlated with behavior initiation, we imaged GCaMP6s while recording leg movements at greater spatio-temporal resolution and trained GLMs on these datasets. As expected, the spatial weightings of these GLMs were very similar to those constructed with jRGECO1a (Extended Data Fig. 6e, R = 0.82). Strikingly, the regions of the brain that predict behavioral activity in these models were those highly enriched for dendritic processes of descending motor neurons, previously described effectors of movement that provide all of the descending connections between the central brain and the ventral nerve cord^22^ (Extended Data Fig. S6). Thus, these data demonstrate that even limited movements recruit many neurons distributed across regions of the brain that contain most of the descending neuron population.

Finally, we note that correlations across brain regions slightly increase during behavior, but do not change in structure, suggesting that intrinsic functional connectivity in the fly is stable, at least over short timescales (Extended Data Fig. 7). Therefore, intrinsic connectivity measures are unlikely to be altered by behavior initiation, at least for leg movement.

## Discussion

By performing the first simultaneous measurements of both neural activity and cellular metabolic flux across the whole brain, our work demonstrates that changes in intracellular calcium levels drive changes in pyruvate concentrations within neurons *in vivo*. These changes in metabolic flux occur on the timescale of seconds, and are strongly suppressed by blocking neuronal firing, arguing strongly that fluctuations in metabolic flux in the brain are driven by neural activity. Moreover, these signals were measured within neuropil, and revealed sharp spatial boundaries between anatomically defined brain regions, demonstrating that the coupling between neural activity and metabolic flux is spatially local, down to the scale of subcellular compartments such as dendrites and axons. As a result, these data provide critical fiducial benchmarks across both space and time, supporting the power of metabolic proxies such as BOLD and FDG-PET to veridically capture changes in neural activity.

By correlating neural activity and metabolic flux with the initiation of behavior, our work reveals that the difference between quiescent states and behavior is coupled to stereotyped, large-scale changes in both activity and metabolic networks in the central brain. Indeed, even small movements of the animal, ranging from a single leg movement, to a bout of grooming behavior spanning seconds, could be well predicted by generalized linear models that positively weighted large regions of the brain that are enriched for the dendritic processes of descending neurons that are engaged in motor control, while negatively weighting other regions. This result is surprising because increasing or decreasing the activities of just single descending neuron types are sufficient to initiate or suppress bouts of walking behavior, arguing for a relatively simple motor command structure to the central brain^23^. By contrast, our finding that all voluntary movements we could measure are associated with large scale changes in neural and metabolic activity across large parts of the brain argue for a much more complex control framework. Thus, even in the relatively compact fly brain, distributed neural and metabolic networks like those described in mice, fish and humans appear to play an essential role in guiding behavior.

## Methods

### Animal preparation

GCaMP6s animals were females of the genotype *w+/w-; UAS-myr::tdTomato/UAS-GCaMP6s; nSyb-Gal4/+*. iATPSNFR animals were females of the genotype *w+/w-; UAS-iATPSNFR/UAS-myr::tdTomato; nSyb-Gal4/+*. Pyronic animals were females of the genotype *w+/w-; UAS-myr::tdTomato/+; nSyb-Gal4/UAS-Pyronic*. Dual Pyronic and jRGECO1a animals were females of the genotype *w+/w-; +/+; nSyb-Gal4/UAS-Pyronic, UAS-jRGECO1a.* Flies were raised on molasses medium at 25°C with a 12/12 hr light/dark cycle. Flies were housed in mixed male/female vials and 5-day old females were selected for imaging.

Flies were prepared as previously described^3^. Briefly, flies were cold-immobilized on ice and placed into a mount separating the head from the body. The frontal parts of the head were removed to allow optical access to the central brain. In sessions not monitoring behavior, legs were immobilized. In sessions monitoring leg movements, legs were kept free.

### Image Alignment and registration

High resolution images were aligned to a template brain and atlas as previously described, except that in dual Pyronic and jRGECO1a imaged flies, a high resolution anatomical scan was made of the jRGECO1a signal instead of myr::tdTomato^3^. Motion correction was performed using AFNI’s 3dvolreg as previously described^3^.

### Two-photon imaging

Flies were imaged at room temperature on a Bruker Ultima system with resonant scanning capability, a piezo objective mount, and GaAsP type PMTs using a Leica 20x HCX APO 1.0 NA water immersion objective lens. GCaMP6s and iATPSnFR signals were excited with a Chameleon Vision II femtosecond laser (Coherent) at 920nm, and collected through a 525/50nm filter. myr::tdTomato signals were excited at 920nm and collected through a 595/50nm filter. Pyronic signals were excited at 860nm and collected through a 525/50nm filter. jRGECO1a signals were excited at 1070nm using a Fidelity II femtosecond laser (Coherent) and collected through a 595/50nm filter. GCaMP6s, Pyronic, and iATPSnFR functional data in Figure 1 and extended data Figure 6 were collected in resonant scanning mode (8kHz line scan rate, bidirectional scanning) and were volumetrically imaged at a resolution of 128×128 (3×3μm) with 68 z sections (3μm steps, effective frame rate ∼100Hz). Dual imaging experiments, representing all other datasets, were collected in galvo scanning mode alternating between 1070nm and 860nm lasers line by line at a resolution of 32×32 (12µmx12µm) with 15 z sections (12µm steps, effective frame rate ∼15Hz).

### Quantification of calcium, pyronic and ATP coupling between ROIs

To measure the coupling between ROI activities in each fly, we first averaged the calcium, pyronic, or ATP signals from all voxels in each ROI to produce a single time series for each ROI and each sensor. We then computed the Pearson correlation of the time series of each ROI pair to generate a 60 x 60 correlation matrix for each fly that represents the couplings between ROIs. We averaged these correlation matrices across all flies to obtain representative correlation matrices (n = 12 flies for GCaMP6s, n = 10 for Pyronic, n = 10 for iATPSnFR, n = 24 for dual-imaged jRGECO1a and Pyronic flies). In order to compare the correlation matrices obtained for these signals, we computed the Pearson correlation between the corresponding average correlation matrices.

### Temporal frequency analysis of neural activity and metabolic flux correlations

Correspondence between neural activity and metabolic flux was measured at a range of frequencies between 0.01 Hz and 0.5 Hz. Using the SciPy open-source mathematical library in Python^24,25^, we applied a Tukey window to the brain signals and then performed a Ricker wavelet transform to decompose the signals into 30 frequency bands. At each frequency band, we measured the Pearson correlation between the filtered calcium and Pyronic signals for each ROI independently, and then averaged this correlation over all ROIs and all flies. We tested against temporally and spatially shuffled signals. For the spatial shuffle, we randomly permuted the identity of the ROIs independently for the pyronic and calcium signals in each fly. For the temporal shuffle, we randomly flipped the temporal axis of each signal independently. Error bars represent the SEM over n = 24 flies. Additionally, we low-passed and high passed one example calcium trace and its corresponding Pyronic trace by setting to zero the Fourier coefficients of these signals above and below 0.1 Hz respectively and computing the inverse Fourier transform of the resulting coefficients.

### TTX application and effect quantification

The jRGECO1a and Pyronic signals were imaged for 1000 seconds prior to TTX application. TTX was then added to the bath through the perfusion at a concentration of 1µM. After a waiting period of 1200 seconds, the brain was imaged again for another 1000 seconds. Analyses were performed on n = 8 flies. The effect of TTX on the Fourier spectrum of the calcium and pyronic signals was measured in two ways. First, for each ROI, we estimated variance by integrating the spectrum between 0.01 Hz and 0.5 Hz (the range of frequencies for which we could correctly estimate the spectrum on a recording of 1000 seconds) before and after addition of TTX (n = 8 flies, error bars are SEM). Second, in order to visualize the influence of TTX at different frequencies, we represented the relative power (difference between the power after addition of TTX and before addition of TTX) as a function of frequency, averaged over all flies and ROIs (n = 54 ROIs, error bars are 95% CI). We also measured the influence of TTX on the coupling between ROIs. For each fly, we computed the correlation matrix between ROI activity over three periods of 500 seconds, with two of the periods taken before the addition of TTX and one period taken after addition of TTX. We evaluated the self-consistency of each coupling before addition of TTX by measuring the Pearson correlation between the first two matrices, and we evaluated the effect of TTX on coupling by measuring Pearson correlations between the second and third correlation matrices. We also measured the effect of TTX on the coupling between calcium and pyronic signals in two ways. First we computed the correlation of their correlation matrices before and after addition of TTX. Second, for each individual ROI, we measured the correlation of calcium and pyronic signals before and after addition of TTX.

### Examination of behavior-related fluctuations in neural and metabolic activity

To measure behavior, we imaged the body of the fly with a c-mount camera (640×512 pixels at 20fps, Flir Blackfly S BFS-U3-04S2M-CS) with a 50mm f./2.0 lens (Edmund Optics). Bouts of activity were manually scored using behavioral observation research interactive software (BORIS)^26^. Flies were scored as behaving if the proximal segments of any of the legs moved. The analyses below were performed on n = 12 flies for jRGECO1a signals, and n = 7 flies for Pyronic signals, all for 2000 seconds. We predicted behavior by fitting a logistic regression, a special case of GLM, on the brain signals of all ROIs. To avoid overfitting, we used an L2 penalty of 1e5 on the weights of the logistic regression. We fitted the model weights on half of the recording and tested the model prediction on the other half for cross-validation (all predictions presented are from the testing phase). In order to assess the predictability of behavior from jRGECO and Pyronic signals, we computed a receiver operating characteristic (ROC) curve and measured the area under the curve (AUC). To assess the predictive power of brain signals at different frequencies, we applied a Ricker wavelet decomposition to both brain signals and behavior and fit a logistic regression model independently for each frequency band. To compare the spatial weight maps of the logistic regression with the smooth processes of descending neurons we used n = 6 flies imaged at 1.2 Hz for GCaMP6s only. To test whether behavior affected coupling between ROIs for jRGECO1a and Pyronic signals, we measured the average difference in coupling across all ROI pairs, between rest and behavior, for each fly independently, and performed a 2-tailed t-test to decide whether this average difference was different than 0.

### Statistics

Comparisons of correlation matrices (Fig. 3d) were performed using a two-tailed paired t-test. Area under curve analysis (Fig 4b) was performed using a one-tailed t-test against 0.5 (no predictive value). Comparisons of correlations during behavior and at rest (Extended Data Fig. 7) were calculated using a one-tailed t-test between the respective correlation values for each ROI.

