## supplemental figures 1-7 for "Causal coupling between neural activity, metabolism, and behavior across the *Drosophila* brain"

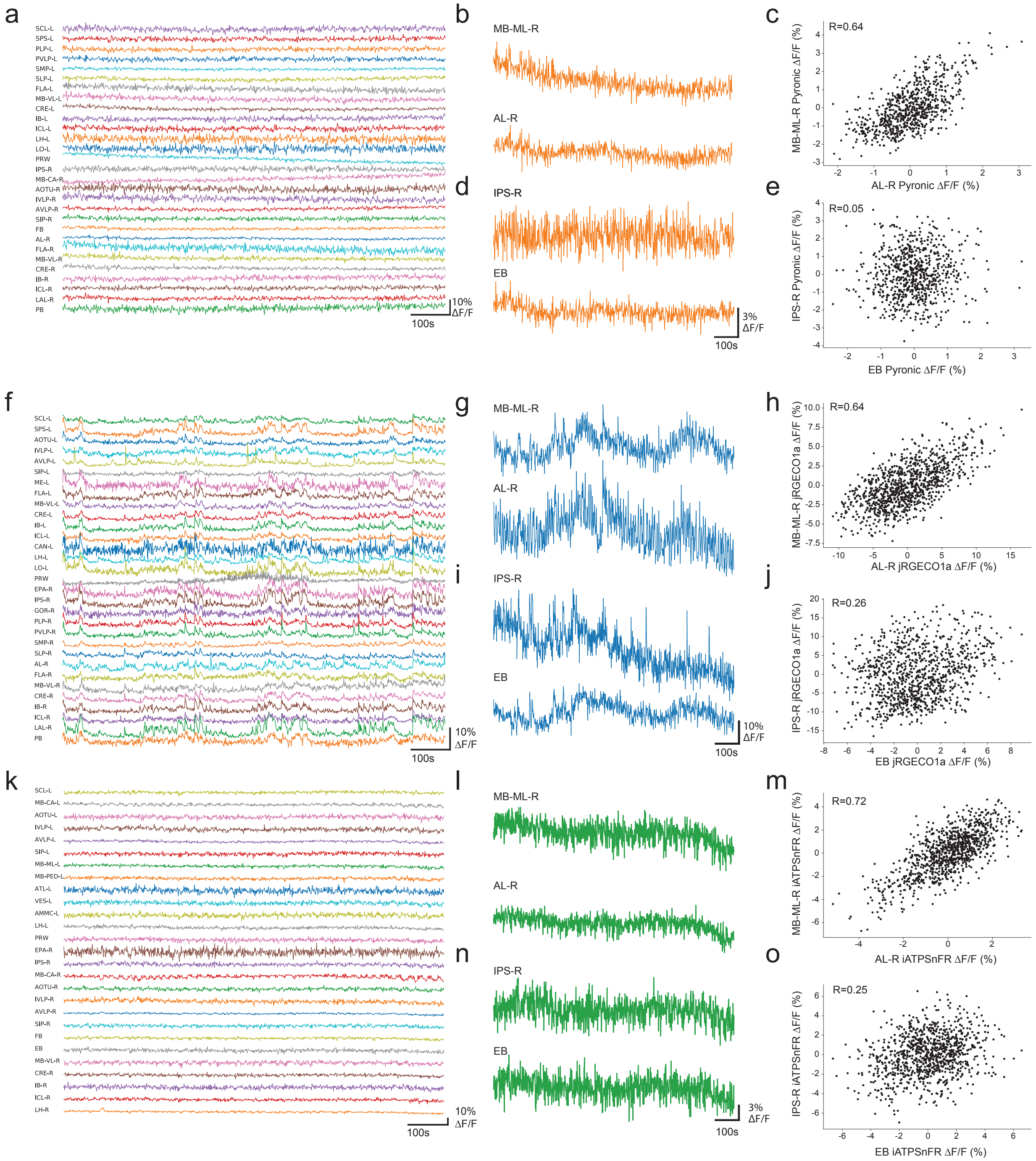


**Extended Data Figure 1. Example traces and correlations of Pyronic, jRGECO1a, and iATPSnFR.** (a). Pyronic traces over an imaging session in different regions. (b). A pair of traces that exhibit high correlation over time. (c). Scatter plot of these two regions demonstrating correlation. (d). A pair of traces that exhibit lower correlation over time. (e). Scatter plot of these two regions demonstrating correlation. (f-j). Same as (a-e) but with jRGECO1a. (k-o). Same as (a-e) but with iATPSnFR.

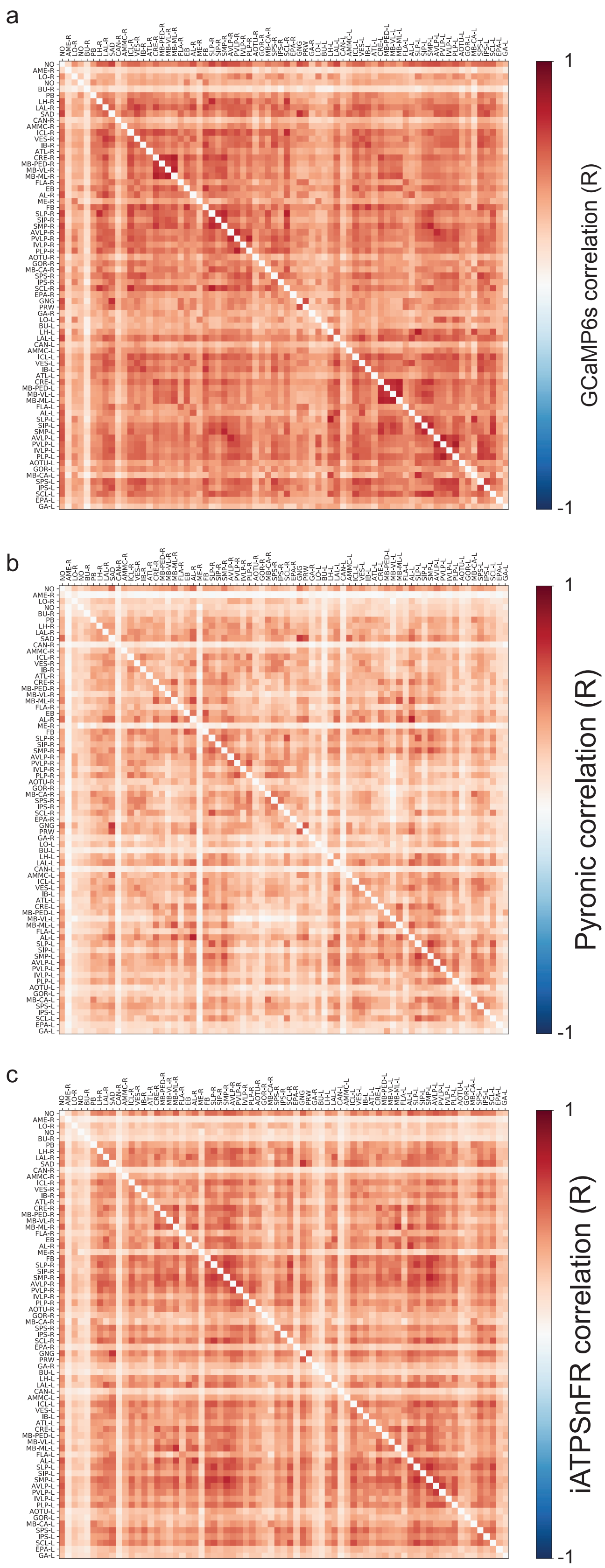


**Extended Data figure 2. Correlation matrices of GCaMP6s, Pyronic, and iATPSnFR**. (a-c) Correlation matrices for GCaMP6s, Pyronic, and iATPSnFR reproduced and enlarged from Figure 1, labeling each individual region.

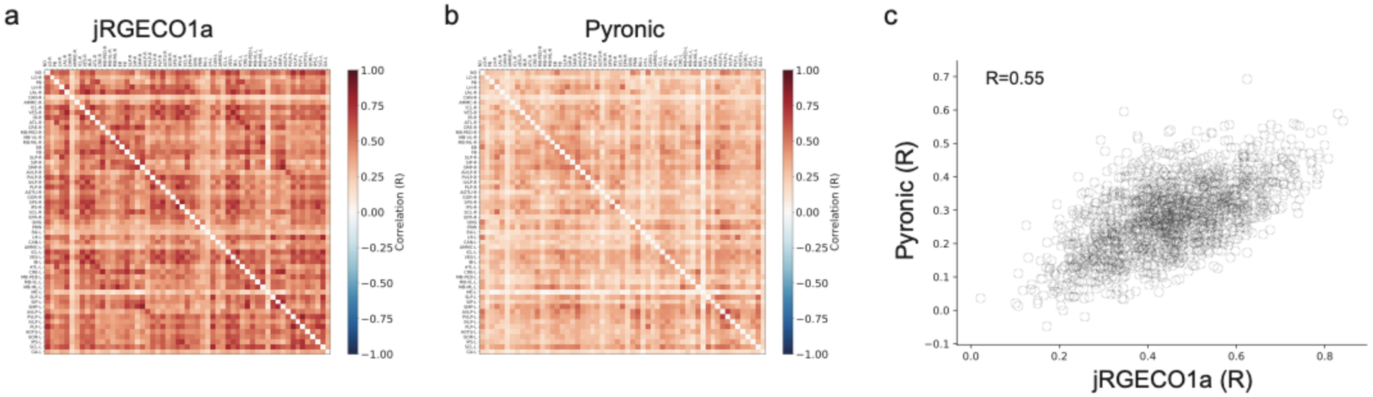


**Extended Data Figure 3. Correspondence of functional networks derived from simultaneous jRGECO1a and Pyronic imaging. (**a). Correlation matrix derived from jRGECO1a in the simultaneous imaging experiments from Fig. 2. (b). Correlation matrix derived from Pyronic in the simultaneous imaging experiments from Fig. 2. (c). Scatterplot of the pairwise correlations between jRGECO1a and Pyronic.

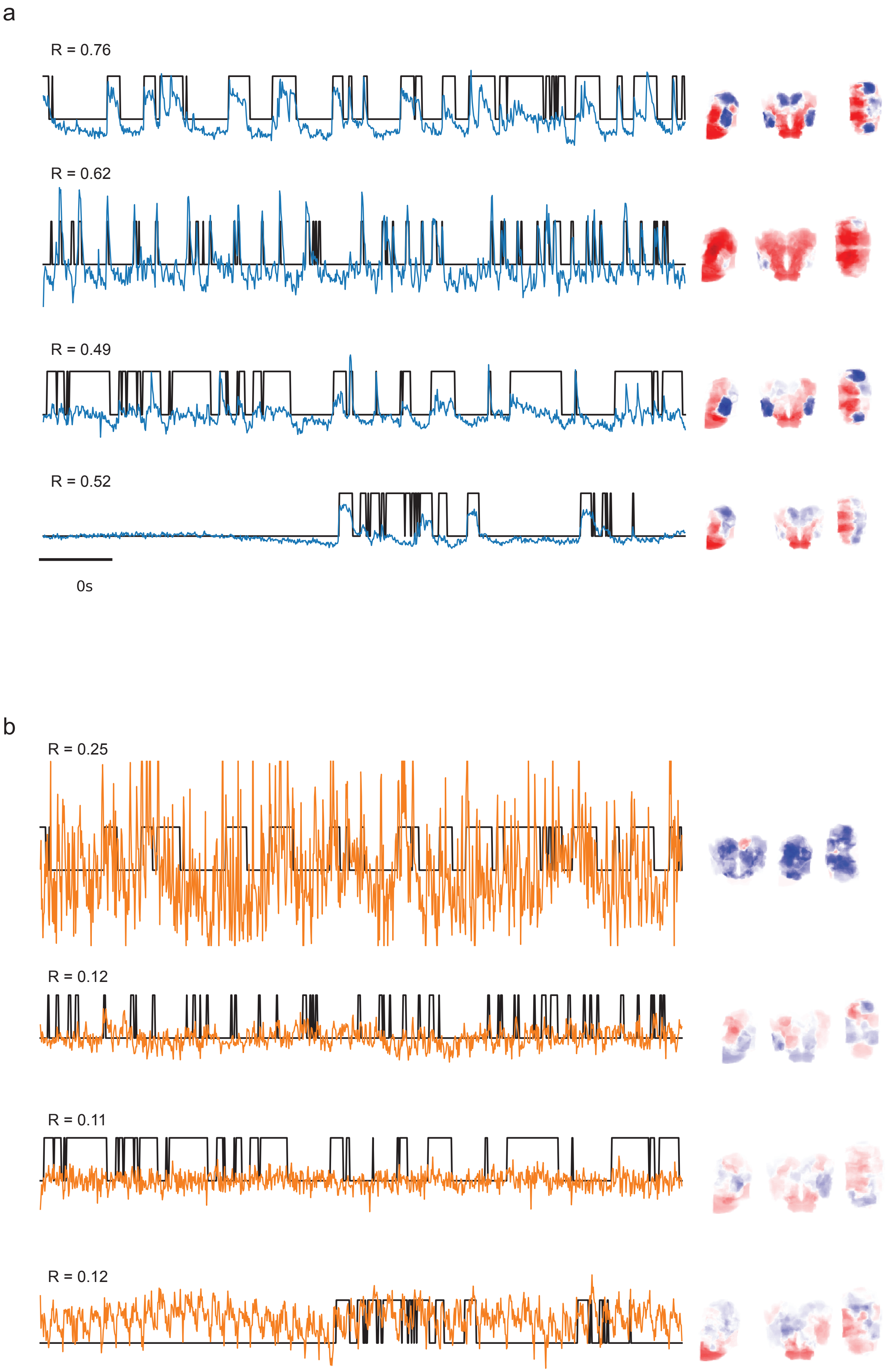
a
R = 0.76
R = 0.62
R = 0.49
R = 0.52
0s
b
R = 0.25
R = 0.12
R = 0.11
R = 0.12

**Extended Data Figure 4. Example model predictions of behavior.** (a). Four example flies showing the prediction based on the model for jRGECO1a (blue) with the corresponding behavior trace (black). Correlation between signals shown above each trace. Weights for each ROI generated by the model shown on right. Oriented as in Fig. 4d. (b). Similar to (a). but with Pyronic (orange).

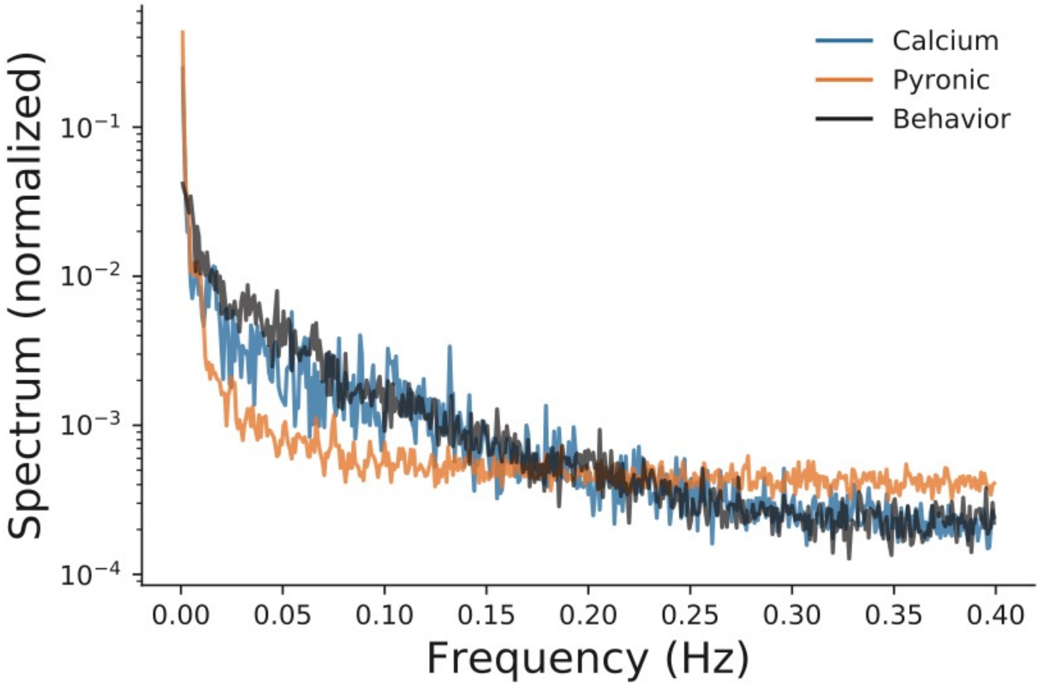


**Extended Data Figure 5. Frequency spectra of jRGECO1a (blue), Pyronic (orange), and behavior (black).** Normalized spectra from data presented in Fig. 4.

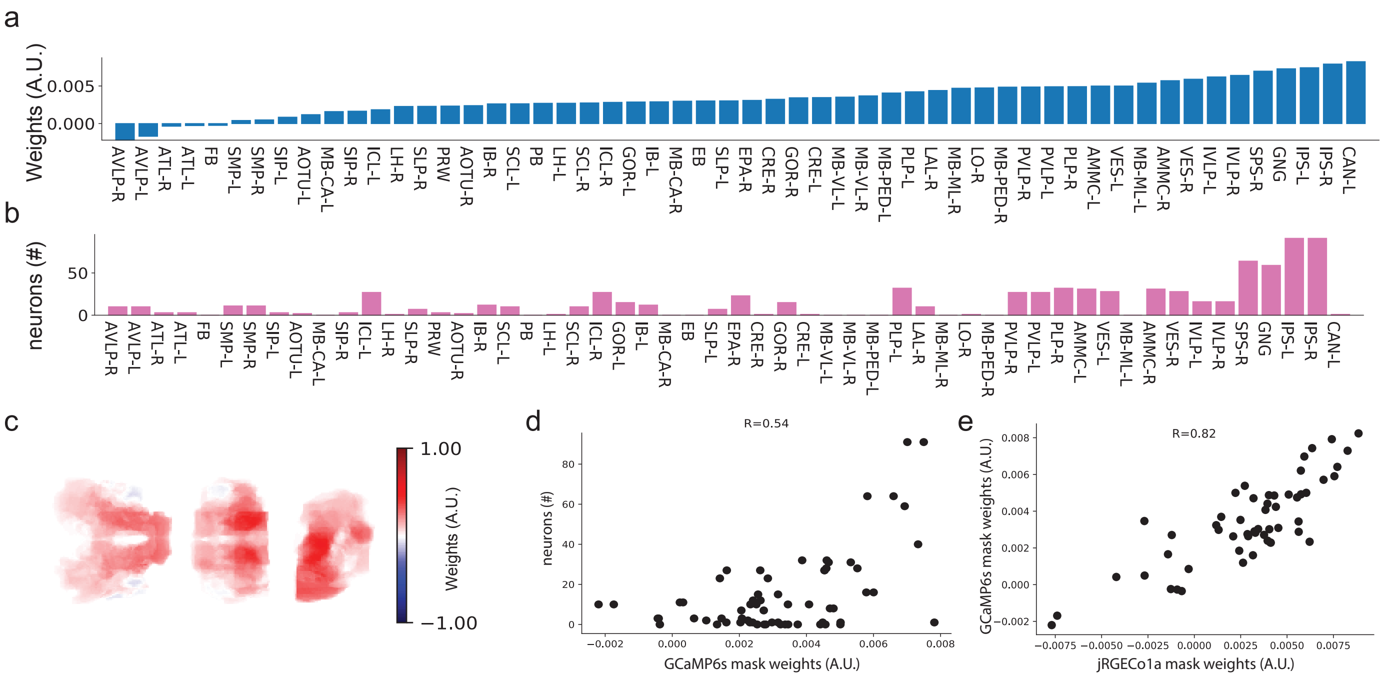


**Extended Data Figure 6. Correlation of model weights for GCaMP6s and descending neuron innervation**. (a). Model weights for each brain region generated using GCaMP6s. (b). The number of descending neuron processes in each brain region[22]. (c). Graphical representation of model weights, similar to Fig. 4d. (d). Correlation between model weights and descending neuron innervation by each region. (e). Correlation between model weights derived from GCaMP6s and jRGECO1a.

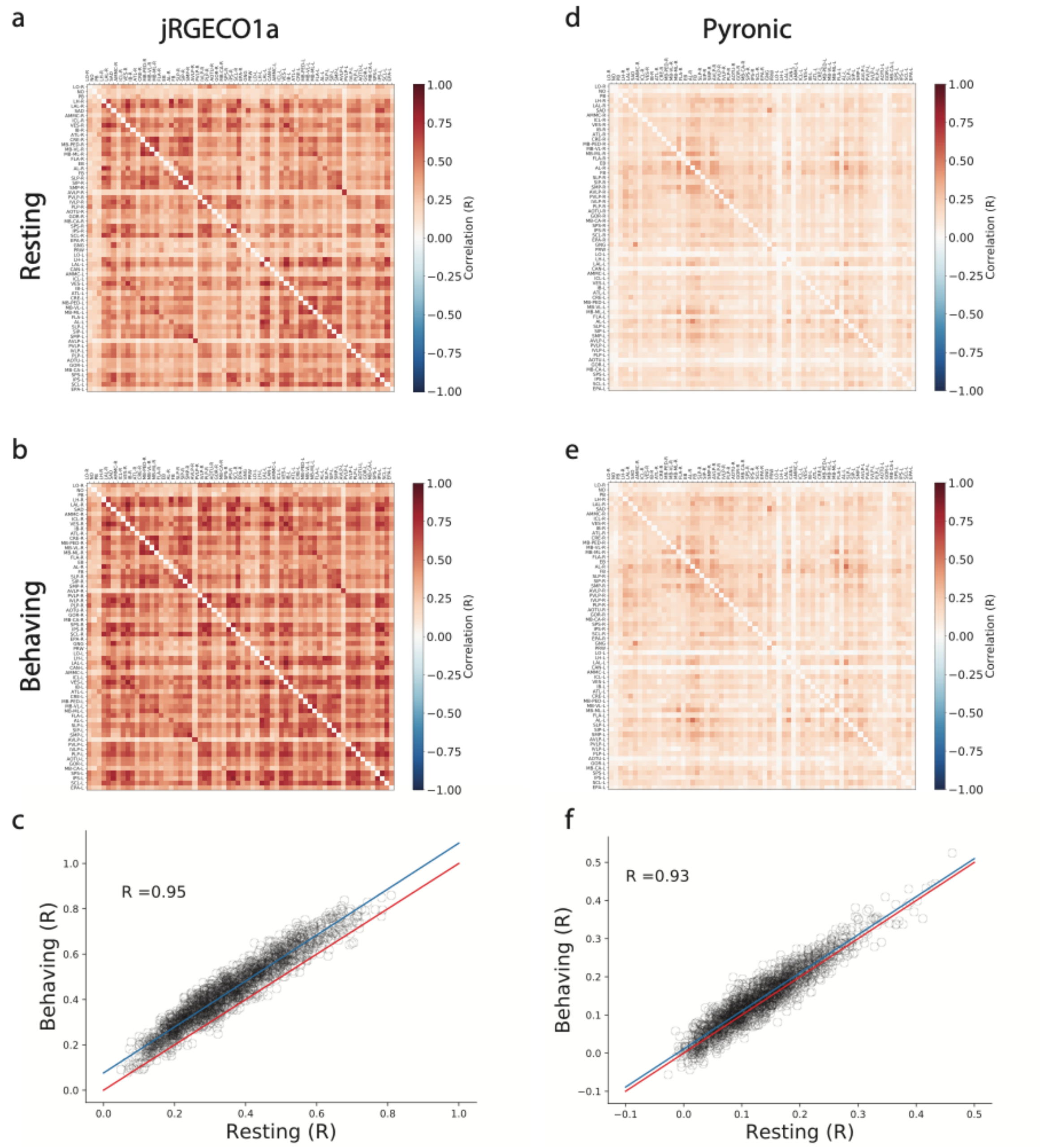


**Extended Data Figure 7. Changes in correlations across regions during behavior for both jRGECO1a and Pyronic**. (a). Functional connectivity map of jRGECO1a during bouts of rest. (b). Functional connectivity map of jRGECO1a during bouts of activity. (c). Correlation of functional connectivity maps during resting and behaving bouts. Correlations increase across the vast majority of regions (P = 0.03, n = 12 flies, one-tailed t-test). (d-f). Same as (a-c) but for Pyronic (P = 0.13, n = 7 flies, one-tailed t-test).
